# Cosmopolitan inversions have a major impact on trait variation and the power of different GWAS approaches to identify associations

**DOI:** 10.1101/2025.09.17.676858

**Authors:** Benedict Adam Lenhart, Alan O. Bergland

## Abstract

The ability of genomic inversions to reduce recombination and generate linkage can have a major impact on genetically based phenotypic variation in populations. However, the increase in linkage associated with inversions can create hurdles for identifying associations between loci within inversions and the traits they impact. As a consequence, the role of inversions in mediating genetic variation in complex traits remains to be fully understood. This study uses the fruit fly *Drosophila melanogaster* to investigate the impact of inversions on trait variation. We tested the effects of common inversions among a diverse assemblage of traits including aspects of behavior, morphology, and physiology, and identified that the cosmopolitan inversions In(2L)t and In(3R)Mo are associated with many traits. We compared the ability of different approaches of accounting for relatedness and inversion presence during genome-wide association to identify signals of association with SNPs. We report that commonly used association methods are underpowered within inverted regions, while alternative approaches such as leave-one-chromosome-out improve the ability to identify associations. In all, our research enhances our understanding of inversions as components of trait variation and provides insight into approaches for identifying genomic regions driving these associations.

**Author’s Summary:** Genomic inversions are large mutations that flip the orientation of sections of DNA, and the presence of inversions has the potential to impact many traits at once. Inversions exist in many organisms including humans and the fruit fly *Drosophila melanogaster*. We take existing knowledge on *Drosophila* traits and identify several inversions which impact many kinds of traits. Approaches such as GWAS can be used to identify the DNA mutations most associated with variation in a trait of interest, but many GWAS methods do not perform well when inversions contribute to variation in phenotype. We show that a common GWAS approach in *Drosophila* is not only unable to find association within inversions, but is overall underpowered. In contrast, we show a different approach is better able to identify mutations within inversions that are potentially associated with fruit fly traits. These findings help scientists studying a wide range of organisms a better understand the role of inversions among different kinds of traits, and supports the broad research of identifying gene associations within these regions of inversion.

## Introduction

Genomic inversions facilitate adaptation by suppressing recombination and generating linkage between many genes and mutations, therefor affecting the genetic basis and evolution of complex traits (1–5). The adaptive importance of inversions for the evolution of novelty, local adaptation, and speciation is clear from a wide variety of organisms across the tree of life (reviewed in (6–10)). Despite the prevalent role of inversions in evolution, the role of inversions underlying phenotypic variation is often overlooked within association studies (11).

Our understanding of the general importance of inversions in affecting trait variation largely comes from ecological genetics, wherein distinct morphs have been identified in natural populations and subsequently linked to inversions (12). For instance, conspicuous behavioral, morphological, phenological and life-history variation has been linked to complex inversion polymorphisms in wild populations of birds (13), seaweed flies (14), monkey-flowers (7), and snails (15). In these cases, and many others (16–18), distinct morphs and their patterns of segregation were first characterized (19–22), prior to identification of inversion genotypes. Therefore, it is less clear if inversions have a major impact on less conspicuous quantitative genetic variation that can be identified through forward mapping approaches, where the goal is to identify the genetic basis of phenotypic variation.

There are two main reasons that forward mapping approaches have potentially missed the impact of inversions on quantitative trait variation. The first reflects the design features of mapping approaches that utilize recombinant populations (23–25). Because inversions reduce recombination, these mapping panels have intentionally used strains with colinear genomes to facilitate recombination and enable efficient QTL mapping. The second reflects statistical techniques of genome-wide association (GWA) studies of outbred wild or laboratory populations. Modern GWA approaches that factor out population structure may have missed important links between inversions and trait variation because inversions can have a major impact on estimates of population structure and relatedness (26–28). Thus, the use of population structure or relatedness estimates as a co-factor in GWA analysis may have led to reduced power to detect association with SNPs linked to inversions. Even when inversion-trait associations can be drawn following GWA (15,29–31), it is challenging to identify the specific genetic architecture within inversion driving association due to the high linkage between loci within inversions (32,33). Therefore, the role of inversions in less conspicuous quantitative genetic variation may have been overlooked in many species.

The fruit fly *Drosophila melanogaster* is an excellent model to assess the importance of inversions on quantitative trait variation. *D. melanogaster* possesses large inversions maintained at intermediate frequencies worldwide and flies show evidence of local and rapid adaptation (reviewed in (34)). *D. melanogaster* inversions are known to impact a variety of traits (35–41). For instance, In(3R)P presence is associated with body size, lifespan, and starvation resistance (37,41), and In(2L)t is associated with behavioral, stress-tolerance, and morphological traits (42–44).

The Drosophila Genetic Reference Panel (DGRP) provides an excellent resource for identifying the effects of cosmopolitan inversions on quantitative variation, and for exploring the role of various GWA methods in discovering associations. The DGRP is a collection of 205 inbred and fully genotyped *D. melanogaster* lines, initially collected from a farmer’s market in North Carolina (45). The lineages have been inbred, their genomes have been sequenced, and the presence of common inversions has been characterized for each line using a combination of polytene chromosome preparations (26), principal component analysis (46), and PCR (47). Due to the availability of these resources, DGRP lines have become a common model for phenotyping studies across many traits (48). To facilitate association studies with the DGRP, several websites have been developed with a standardized mapping approach that factors out inversions and other drivers of relatedness (26,48). Indeed, GWA approaches that correct for genome-wide structure or inversions account for approximately 60% of DGRP studies from a representative sample of 36 papers (curated dataset;(48)), yet few (35%) report testing for associations with inversions or inversion linked markers (Supplemental Table 1).

In our study, we test the ability of different GWA mapping approaches to identify signatures of association with inversions and linked variants. We utilize published studies that measured phenotypic variation in the DGRP. First, we show that several cosmopolitan inversions have large effects on dozens of traits, in that they explain more trait variation than expected by SNPs of comparable frequencies. Next, we explored four genome-wide association strategies that differ in their genetic-relatedness matrices (GRMs) and the treatment of inversions as co-factors, and contrast the real GWA signal for each phenotype to 100 permutations. We generated three types of GRMs (i) using the full genome, (ii) using an LD-thinned genome, (iii) and using a leave-one-chromosome-out (LOCO) approach. In addition, we performed association analysis and permutations using the full-genome based GRM and factored out the effect of inversions following methods outlined in (26,48). We show that the result of the GWA greatly depends on the mapping strategy, and that only the LOCO approach resolves association signals that exceed permutations. Finally, we use the output of the LOCO-GWA to test whether SNPs identified as top candidates under the different mapping strategies show different levels of enrichment for signatures of local adaptation, and whether signals of pleiotropy are resolvable at specific loci inside the inversion.

## Materials and Methods

### Selection of trait data

We reanalyzed trait data collected on the DGRP (45). We made use of the DGRPool resource, which has consolidated the inbred line means from many publications (48). We used the “curated” data, and removed traits from this dataset that describe genomic features (such as genome size, P-elements, transposon presence) or used less than 75 unique DGRP lines, ending up with 409 unique traits derived from 36 publications (Supplemental Table 1). Some traits measure the same or similar phenotype, and thus there is potential for our results to be biased toward more frequently measured phenotypes. Of these 36 studies, 19 of them were also represented in the independent meta-analysis reported in Nunez et al., 2024 (43). We annotated these traits by classifying each trait into 5 general groups: “Behavior”, “Life-History”, “Morphology”, “Physiology”, and “Stress-resistance” (Supplemental Table 1).

### The phenotypic impact of cosmopolitan inversions

We characterized the effect of cosmopolitan inversions In(2L)t, In(2R)NS, In(3L)P, In(3R)K, In(3R)P, and In(3R)Mo on these traits, as these are the inversions considered by the DGRP analysis webtool. For this analysis, we used the inversion classifications provided by (26). These classifications reference the same 205 DGRP lineages present in the DGRP genotyping data Supplemental Table 2 For each trait, we used a simple linear model to test the effect of any single inversion using strains that were homozygous for either the inverted or standard allele. We did not include heterozygotes as there is no way to determine the frequency of the inverted allele within DGRP lines marked as heterozygous, nor to know the genotype of any individual that was phenotyped. In addition, we do not consider the impact of multiple inversions simultaneously, nor do we test for interactions between inversions because of the relatively low frequency of inversions in the dataset.

For each phenotype, and for each inversion, we compared the results of the linear model with a null model using an ANOVA test, and counted the number of traits with significant association with any of the inversions with p-value < 0.05 (Supplemental Table 1). Next, we evaluated whether the extent of association between traits and inversion status is greater than expected relative to other random polymorphisms in the genome. The motivation for this analysis is to test if the inversions have a greater impact than expected by chance. For each inversion, we replicated the above linear modeling using 100 SNPs and small indels randomly selected from those identified at the same frequency as each inversion (±1%), and on the same chromosome arm, but at least 2MB from the inversion breakpoints to avoid areas of highest linkage disequilibrium. By comparing the number of traits significantly associated (p < 0.05) with the inversions and random polymorphisms, we can use the 100 random polymorphisms to approximate the effect of a given mutation on trait data expected by chance. Last, we calculated R^2^ (coefficient of determination) for the observed and matched-polymorphism models to ask whether the inversions explain more variation than expected by other comparable SNPs and small indels in the genome.

If a trait was significantly associated with an inversion using the linear model results, and if the R^2^ of that model surpassed the 95% quantile of permutations, we designated that trait as part of a group of “inversion-associated traits” used in downstream analysis. From the original five inversions, in subsequent analysis we only pursued inversions in which at least 5% of the samples used were homozygous for the inversion. We identified two sets of inversion-associated traits, containing traits associated with In(2L)t or In(3R)Mo.

One last extension of this analysis was to examine the impact of the ancestry of the DGRP lines on trait line averages. This is important because North American *D. melanogaster* populations, including the DGRP, result from secondary contact and admixture between European and African populations about 150 years ago (49–52). Using estimates of the proportion of European and African ancestry of each DGRP line from Pool 2015 (52), we created linear models using the same statistical tools as described above. For each phenotype, and each inversion, we evaluated an Ancestry-only model (ancestry as a fixed effect), Inversion-only model (inversion genotype as a fixed effect), and a Full model (both ancestry and inversion genotype as fixed effects) against a null model using ANOVA. Observed models were compared against permutations in which phenotype line averages were shuffled. Last, we repeated this statistical framework, this time comparing the Full model against the Ancestry and Inversion models.

### Principal component analysis of trait data

We used principal component analysis (PCA) on each of the inversion-associated phenotypic trait sets (Supplemental Table 3). Missing data for any trait were imputed using the *imputePCA* function from missMDA v1.19 (53) using the “regularization” method. This approach uses the average phenotype value for initial imputation, and performs a secondary regularization step. We used the imputed data to conduct PCA using FactoMineR v2.8 (54). We separated the principal component projections by inversion presence using the inversion genotype of the DGRP lines and then compared the PCs of “inverted” and “standard” groups using Student’s t-test.

### Principal component analysis of genomic data

To understand how inversions impact general patterns of multi locus genetic variation, we performed a series of PCA on the DGRP genetic SNP and small indel polymorphism data. We used three polymorphism selection strategies for this principal component analysis that mirrors polymorphism selection strategies used for the construction of the GRM (see below). The first version of polymorphism selection (“Full”) used all SNPs and small indels across the autosomes and X chromosome with minor allele frequency (MAF) greater than 5% and sites with missing genotype data in less than 20% of DGRP lines. The second version (“LD”) used SNPs and small indels with MAF > 5% and missing rate < 15%, and a low pairwise linkage disequilibrium (R^2^ < 0.2). To ensure that SNPs were at least 5000 base pairs apart we used the *snpgdsLDpruning* function of the R-package SNPrelate v3.17 (55), with the *slide.max.bp* parameter set to 5000. The third version used a leave-one-chromosome out (“LOCO”) approach (56) that used the same filtering and thinning strategy as the LD-pruning approach but in four parts, each missing one of the main chromosomal arms in order to reduce the effect of an inversion on relatedness within its own chromosomal arm. We used the *snpgdsPCA* function from *SNPrelate* v3.17 (55) for PCA. To quantify the effect of inversions on PC1-PC4, we constructed linear models in the same manner described above, recording the R^2^ of both the linear models and a set of permutations with the lines’ inversion genotype shuffled. We designated a model outcome significant if its R^2^ surpassed 95% of permutations.

### Construction of GRMs

We developed three genomic relatedness matrixes (GRM) to address population structure in different ways. For the “Full” method, we use the GRM matrix that is supplied by the DGRP website and is commonly used in DGRP GWAS studies (http://dgrp2.gnets.ncsu.edu/, last accessed 04/20/2025). This approach uses the VanRaden method (57) to construct a GRM from all SNPs and small indels with a MAF > 0.05 and a missing rate < 20% (26). For the “LD” method, we used LD pruning using the same parameters that we used for the LD-pruned PCA, described above, and constructed a GRM from the whole genome using the *snpgdsGRM* function in SNPRelate based on the Genome-wide Complex Trait Analysis (GCTA) method (58). For the “LOCO” method, we generated sub-GRMs, each one drawing from the DGRP genome but ignoring one chromosome arm (“2L”, “2R”, “3L”, “3R”, and “X”) and using the same steps as described for the LD-thinned approach.

### GWA Analysis

We performed association mapping using mixed-effect models implemented in the R package *GMMAT* v1.3.2 (59). This approach used either the “Full”, “LD”, and “LOCO” GRMs as a random effect to control for population structure. In addition, we performed a fourth association mapping approach based on the GWA approach developed by Huang *et al.* 2014 (26), which we refer to as the “Factored-out” approach. The Factored-out approach first standardizes each trait by the effects of the inversions by regressing line mean data against inversion status using the model:

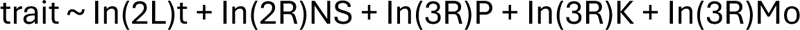

Next, the residuals of this model are used as the trait or association analysis. We used the “Full” GRM with the Factored-out approach to replicate the association model implemented in by Huang et al. 2014 (28) and available on the DGRP online GWA tool (http://dgrp2.gnets.ncsu.edu/).

For each of the four GWA approaches, we compared a “reduced model” to a “full model.” The reduced model is described by the formula:

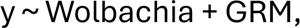

where *y* represents the line means for a particular trait (or residuals in the case of the factored-out model). Wolbachia is an infectious symbiote known to affect aspects of Drosophila fitness (60–62), and is included as a cofactor in standard DGRP GWA approaches. Here we encode Wolbachia infection status as a fixed effect listed as present or absent, based on the tables published in Huang et al., 2014 (26). GRM is a random effect genetic relatedness matrix. The reduced model is compared to a full model defined as:

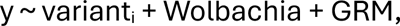

where variant_i_ is the fixed effect and an additive representation of the dosage of the *i*^th^ SNP or small-indel reported for the DGRP. We contrasted the full and reduced models using the *glmm.score* function in the GMMAT package (v1.4.2), which filters out all variants with minor allele frequency < 5% and missing data > 15%. In the LOCO approach (56), we split the scoring of the genome into five sections for each of the major chromosomal arms, with each GWAS using the sub-GRM constructed without that corresponding region. Our GWA approach scores inverted and non-inverted regions without distinction.

For each trait and GWA method, we conducted 100 permutations by randomly shuffling the trait data prior to fitting the reduced and full models.

### GWA summary statistics

We compared overall genomic signal from the GWA of each trait using statistics for the observed and permutated GWA models. We partitioned each chromosome into bins based on whether SNPs are inside or outside inversions as defined using coordinates in Corbett-Detig et al. 2012 (47). For each bin, we calculated the proportion of SNPs with a p-value less that 10^-5^ (“hits”), a common p-value threshold in DGRP studies (63–69), as well as the genome inflation factor (GIF) as the ratio of the median observed variant-wise *X*^2^ values within the bin divided by the expected median *X*^2^ with 1 degree of freedom. We compared the summary statistics in the observed data to the statistics from the permutations for each trait and GRM method, reporting the proportion of traits where the statistic exceeds the 95th percentile of the trait’s permutation-based distribution.

### Enrichment tests

We tested if GWAS hits identified via different GRM approaches prioritize SNPs that are potentially subject to temporally or spatially variable selection using BayPass version 2.1 (70). We used data from the DEST dataset to obtain allele frequencies from *D. melanogaster* population*s* sampled across the North American East Coast (71), and Charlottesville, Virginia across multiple years (43). We identified polymorphisms that are more differentiated than expected given population structure (XtX* outliers) and tested association between variants and environmental variables after correcting for population structure. Using this framework, we identified differences in allele frequencies between populations (XtX*) and Bayes Factor (BF) of association with environmental variables. We used latitude for the association of East Coast variants and maximum temperature two weeks prior to collection for Charlottesville variants (43). We ran the software five times and report the mean statistic per SNP (72). We generated a null distribution of XtX* and BF using the POD (Pseudo-observed Data) framework for 10-times the number of SNPs as the observed data, ran BayPass with five replicate iterations, and calculated empirical p-values for XtX* and BF using these POD simulations.

We identified the level of enrichment between top GWA variants and top BayPass variants. We identified the top hits within each GWA study by identifying the 500 hits with the lowest P-value, and top XtX* and BF variants as those that surpass 95% of the corresponding distribution from the simulated POD data. We computed Fisher’s Exact test by contrasting the odds that top association hits for any trait are enriched for top XtX* and BF hits. We compared these Fisher’s Exact test odds-ratios to odds-ratios constructed in the same way using the permuted GWA.

### COLOC analysis

We tested if inverted regions of the genome are likely to have pleiotropic effects on phenotypic variation using the *coloc.abf* function from coloc v5.2.3 (73). By treating the top principal component projections as dimensionality-reduced traits, we sought to identify regions in the genome with a shared association with multiple inversion-linked traits. We used the Factored-out and LOCO GWAS frameworks to score the impact of SNPs genome-wide on the PC1 and PC2 loadings for the In(2L)t and In(3R)Mo associated traits. We identified areas of colocalized signal on PC1 and PC2 using a sliding window analysis across the genome with window size 10Kb and step size 5Kb, and comparing the SNP GWAS data using *coloc.abf*(). This analysis identified regions of SNPs likely associated with the traits differentiated along only PC1, only PC2, or regions of SNPs with a colocalized association across PC1 and PC2.

## Results

### Cosmopolitan inversions impact phenotypic variation

To study the role of inversions on genetically-based trait variation in *D. melanogaster*, we reanalyzed data from publications that measured trait variation in the DGRP and were curated in the DGRPool database (48). We analyzed 409 traits, categorizing them into five groups: morphological, life-history, stress resistance, physiological, and behavioral (Supplemental Table 1). We found that In(2L)t, In(3R)Mo, and In(3R)K are associated with more traits than expected given SNPs of the same frequency (**Fig. 1A**). In(2L)t is especially associated with many behavioral traits including startle response, sleep, and movement, while In(3R)K and In(3R)Mo are associated with morphological traits such as femur and abdomen size (Supplemental Table 1). We found that the inversions explain ∼10% of the variation in these traits, and that dozens of traits are explained better by inversion status than expected from random polymorphisms in the genome (**Fig. 1B**).

**Figure 1.**
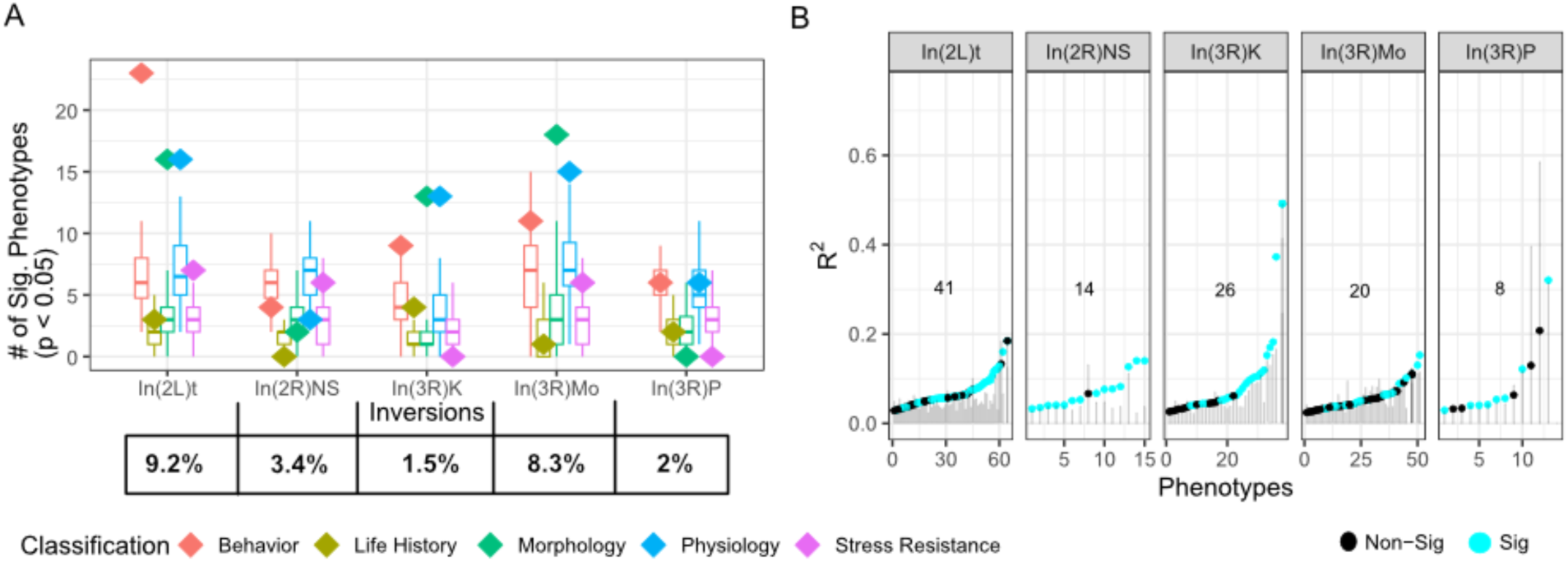
Cosmopolitan inversions have a major impact on trait variation. **A)** Diamonds, colored by trait category, indicate the number of traits significantly affected by inversion presence at p<0.05, overlaying the same-frequency models shown with box plots. The percentage of lines homozygous for each inversion within the lineages tested is given at the bottom. **B)** The proportion of variation explained by each inversion (R^2^) for traits significantly associated at p<0.05 with each inversion, compared against the distribution of corresponding same-frequency models in grey. Statistically significant inversion model values that surpass the null distribution are colored cyan. The number in each panel of B is the count of phenotypes where R^2^ exceeds permutations and traits that are significantly associated with the inversion with p<.05.

### Principal component analysis of traits associated with In(2L)t and In(3R)Mo

To better understand the directionality and intensity on inversion impact on traits, we performed PCA on the traits that are associated with In(2L)t and In(3R)Mo and also explain more variation than expected by chance (In(2L)t: n=41, In(3R)Mo: n=20; Fig. 1). The top two principal components (PC1 and PC2) explain over one third of the trait variation for both In(2L)t and In(3R)Mo (**Fig. 2A**). Therefore, we restrict further analysis to these two principal components. In(2L)t significantly loads into PC1 (t-test, t = -4.38, df = 16.68, p = 4.29e-4) of its associated trait set, while In(3R)Mo significantly loads onto both PC1 (t-test, t = -5.18, df = 18.03, p = 6.21e-5) and PC2 (t-test, t = 2.27, df = 15.54, p = 0.038; **Fig. 2B**) of its associated trait set. In the In(2L)t PCA, traits like body size and ethanol sensitivity have positive loadings on PC1 and traits like startle response and negative geotaxis have negative loading. Lines homozygous for In(2L)t have lower values of PC1, thus have higher startle response and higher activity levels (**Fig. 2C**), among other differences (Supplemental Table 3). In the In(3R)Mo PCA, traits like body size have positive loadings on PC1 and traits like feeding and chill coma recovery have negative loading. Lines homozygous for In(3R)Mo have lower values of PC1, thus have lower body size (**Fig. 2D**), among other differences (Supplemental Table 3).

**Figure 2.**
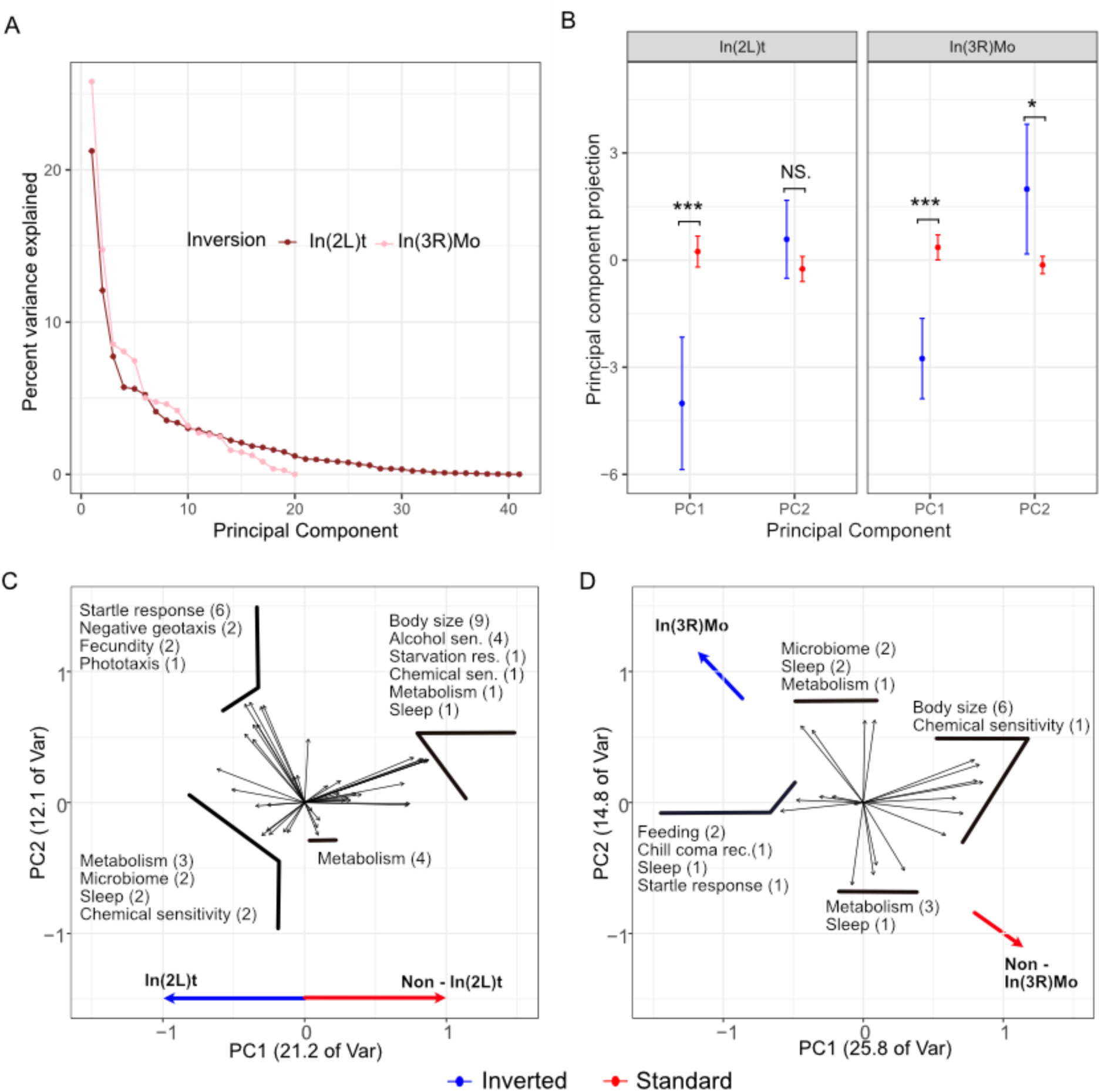
Principal component analysis of In(2L)t and In(3R)Mo. **A)** Scree plot showing the variance explained by principal components, colored by their associated inversion. **B)** The projections of In(2L)t and In(3R)Mo genotype onto PC1 and PC2 projections. **C)** PC loading values for the traits significantly impacted by In(2L)t. Labels are aggregated to show similar traits together, e.g. “Sleep (2)” corresponds to two sleep traits. Variance explained by each PC is given on the axis title. **D)** PC loading values for the traits significantly impacted by In(3R)Mo.

### Controlling the effect of inversions on PC and GRM space

Cosmopolitan inversions in *D. melanogaster* have been shown previously to have an impact on genome-wide patterns of genetic variation as summarized by principal components and genetic relatedness matrices (27,28,74). Therefore, we tested if different polymorphism selection strategies can mitigate this impact. As previously reported (26), PCA of the “Full” genome shows that inversions strongly impact PC space. In(2L)t primarily impacts PC1F_ull_ (F _1,179_ = 870, p = 2.89e-70) and In(3R)Mo primarily impacts PC2_Full_ (F _1,195_ = 331, p = 5.54e-44, **Fig. 3A**). The “LD” polymorphism-set reduces the significance of inversion impact on PC1_LD_ for In(2L)t (F_1,179_ = 175, p = 2.89e-28) and the impact on PC2_LD_ for In(3R)Mo (F_1,195_ = 90.71, p = 6.93e-18, **Fig. 3A**). In contrast, PCA of the LOCO genome shows a sharply reduced impact of In(2L)t on PC1_LOCO_ (F _1, 179_ = 5.84, p = 0.017) and In(3R)Mo on PC2_LOCO_ (F _1,195_ = 0.02, p = 0.88, **Fig. 3A**).

We calculated the proportion of variation in the genetic PC1 and PC2 that is explained by inversion status, and contrasted that to a null distribution made via 100 permutations. We found in the Full and LD methods, In(2L)t and In(3R)Mo explained more variation for PC1 and PC2 than expected by chance (**Fig. 3B**), with In(2L)t explaining the most variance within PC1 using the Full method and less using the LD method, while In(3R)Mo explained the most variance for PC2 within the Full method and less within the LD method. Meanwhile, within the LOCO method each inversion explains a near zero amount of variance for PC1 or PC2, and PC2 is no longer significantly impacted by either inversion (**Fig 3B**). For PC3 and PC4, the R^2^ of each of the principal component ∼ inversion genotype models was near zero, indicating there is little correlation between inversion genotype and these principal components (Supplemental Figure 2).

To identify how the presence of inversions impacts patterns of relatedness, we compared the relatedness of the DGRP lines using different polymorphism selection strategies. Across all approaches, relatedness is low within standard genotype lines with no cosmopolitan inversions (**Fig. 3C**). This replicates observations in Huang et al. 2014 (26). However, within the Full and LD-thinned approaches we see that relatedness is driven higher within lines homozygous for an inversion. In contrast, the LOCO approach on a given chromosome can drive relatedness for homozygous inverted lines near zero while still accounting for inversions on the other chromosomal arms (**Fig. 3C**).

**Figure 3.**
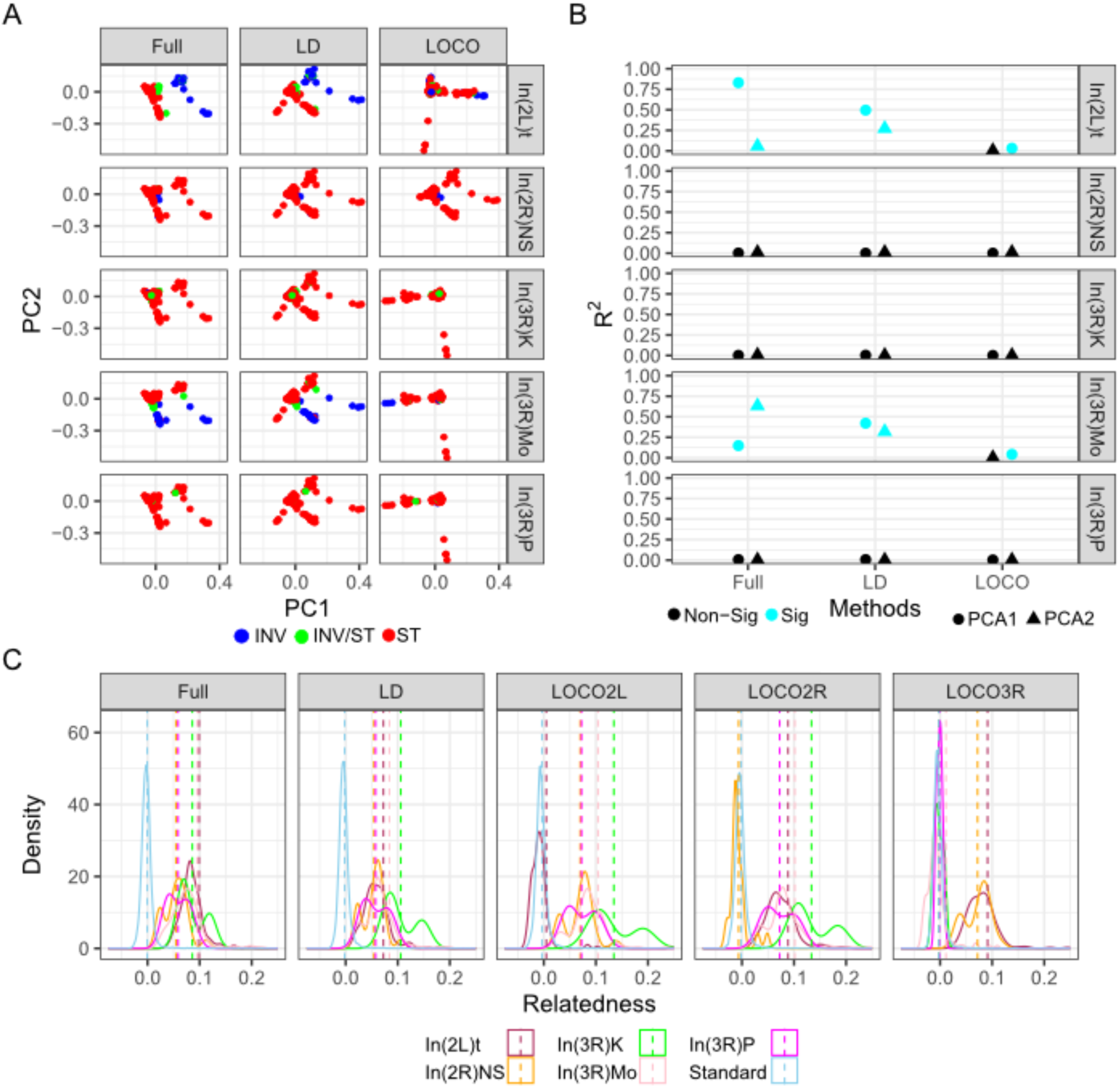
Inversions influence population genetic structure and relatedness. **A)** The first and second genomic PCs for each sample colored by the genotype of that sample. **B)** The R^2^ values for models comparing PC1 and PC2 to inversion, colored by which values exceed a distribution of permutations. **C)** The distribution of pairwise relatedness values between each of the samples, indicated by a solid line colored by genotype of the samples and split across di<erent methods, with a dotted line indicating means.

### The LOCO approach can better capture signal for inversion-associated traits

After characterizing the impact of different GRM methods and presence of inversion in the DGRP data, we tested the strengths and weaknesses of the four GWA strategies (Factored-out, Full, LD, LOCO) on the magnitude of association signal. We compared the summary statistics from the observed trait GWA against permutations to see how many traits identified more signal than expected by chance. Using the Factored-out approach, we found that about 6% of traits have “hit-counts” that exceeds the largest 95% of the null distribution generated by permutation (**Fig. 4**). In other words, the Factored-out approach largely fails to identify more associations than would be expected by random chance, as 6% is about the number of traits that would surpass permutations as false positives. We found that the Full and LD-thinned approaches also fail to identify many more significant associations than expected given the permutations. The LOCO method surpasses permutation significantly more often than the Factored-out method within inverted regions (Fisher’s Exact Test - FET, 2L: p = 9.47e-9, 2R: p = 5.17e-4, 3L: p = 6.66e-10, 3R: p = 1.61e-7), as well as outside the inverted region (FET, 2L: p = 1.2e-9, 2R: p = 7.71e-4, 3L: p = 1.61e-5, 3R: p = 9.19e-13) (**Fig. 4**). Similarly, the GIF of LOCO surpasses permutation significantly more often than the Factored-out method within inverted regions (Fisher’s Exact Test - FET, 2L: p = 7.05e-45, 2R: p = 3.14e-12, 3L: p = 7.42e-6, 3R: p = 2.21e-29),, as well as outside the inverted region (FET, 2L: p = 1.59e-19, 2R: p = 1.38e-13, 3L: p = 2.35e-5, 3R: p = 1.15e-21: (**Fig. 4**). However, this increase in GIF via LOCO is not uniform across the chromosome. The GIF of traits scored with LOCO is significantly higher when scored in inverted regions then non-inverted regions on both 2L (FET, p = 4.34e-8) and on 3R FET, p = .014).

**Figure 4.**
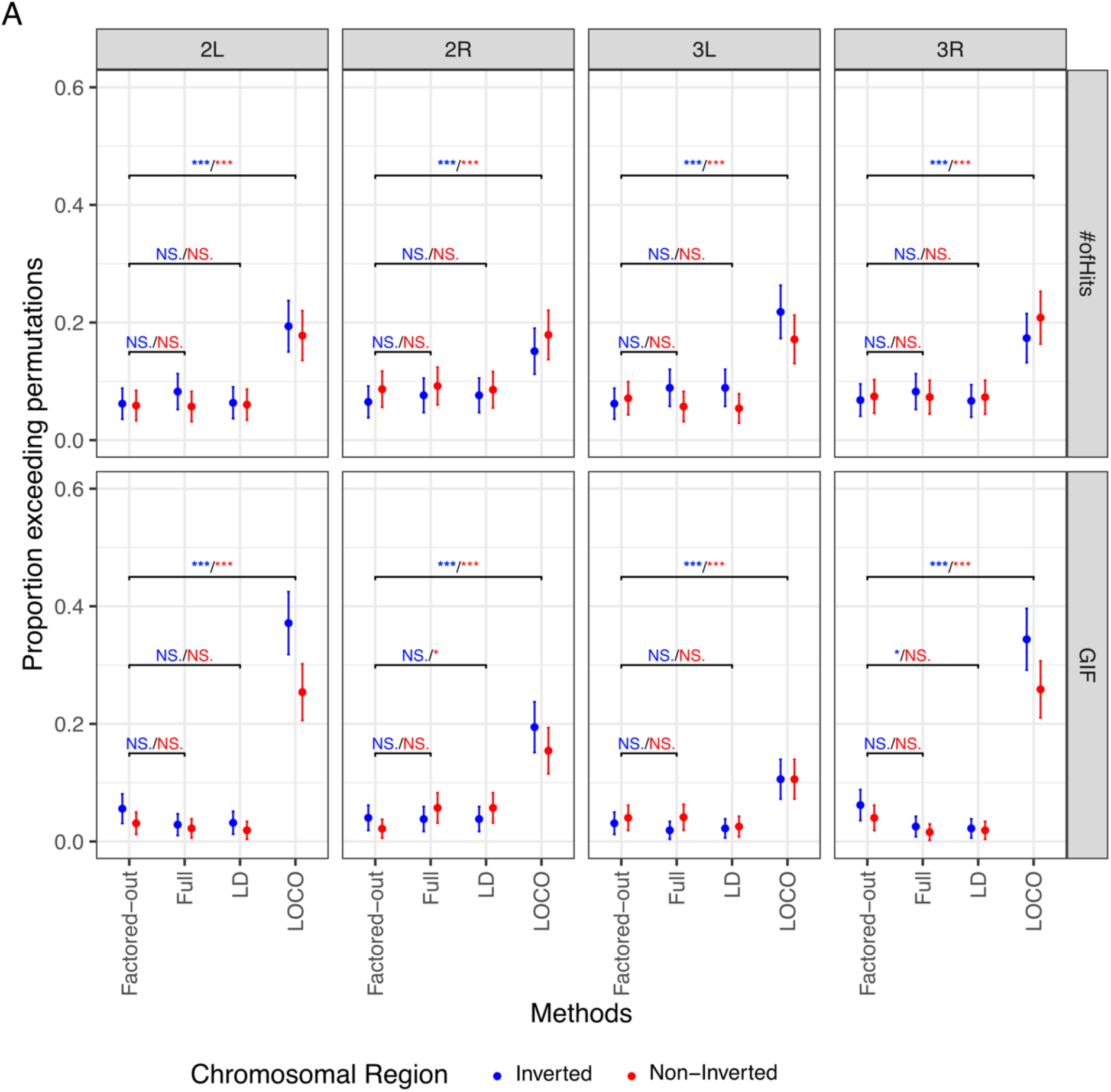
The LOCO approach can better capture signal genome-wide. **A)** The proportion of significant hits and GIF for each GWAS output are compared across each method and colored by location relative to inversions. The proportion of traits that exceed their corresponding permutations is given on the y axis, along with binomial confidence intervals. The color of the significance annotation refers to the inversion genotype under comparison. (* = p < 0.05, ** = p < 0.01, *** = p < 0.001, NS. = p >= 0.05)

### Enrichment tests

To compare the utility of the four approaches to identify biologically meaningful loci, we characterized the ability of LOCO and Factored-out methods to identify loci thought to be important for local adaptation. Using estimates of allele frequencies of *D. melanogaster* collected across seasons and across latitudes (71,75), we used the Baypass (70) software to identified SNPs that are more strongly differentiated across the North American east coast, or within Charlottesville, VA through time (XtX* outliers). In addition, we identified the strength of association between SNPs and latitude for the East Coast samples and between SNPs and temperature in the two weeks prior to sampling for the Charlottesville samples. To understand which methods could successfully identify enrichment within inverted regions, we (i) examined the enrichment on SNPs within inversions in 2L and 3R and (ii) compared enrichment between traits associated and not associated with the corresponding inversion. There was a significant jump in enrichment between GWAS hits and max temperature bayes factor for the LOCO method on 2L (FET: p = 0.019), but not for hits derived from the Factored-out method (**Fig. 5**). Correspondingly, there was an increase in enrichment between GWAS hits and differentiation across max temperatures for the LOCO methods on 3R (FET, p = 0.011) but not for the Factored-out method (**Fig. 5**). There was no difference reported within the East Coast enrichments with GWAS hits between inverted and non-inverted associated traits.

**Figure 5.**
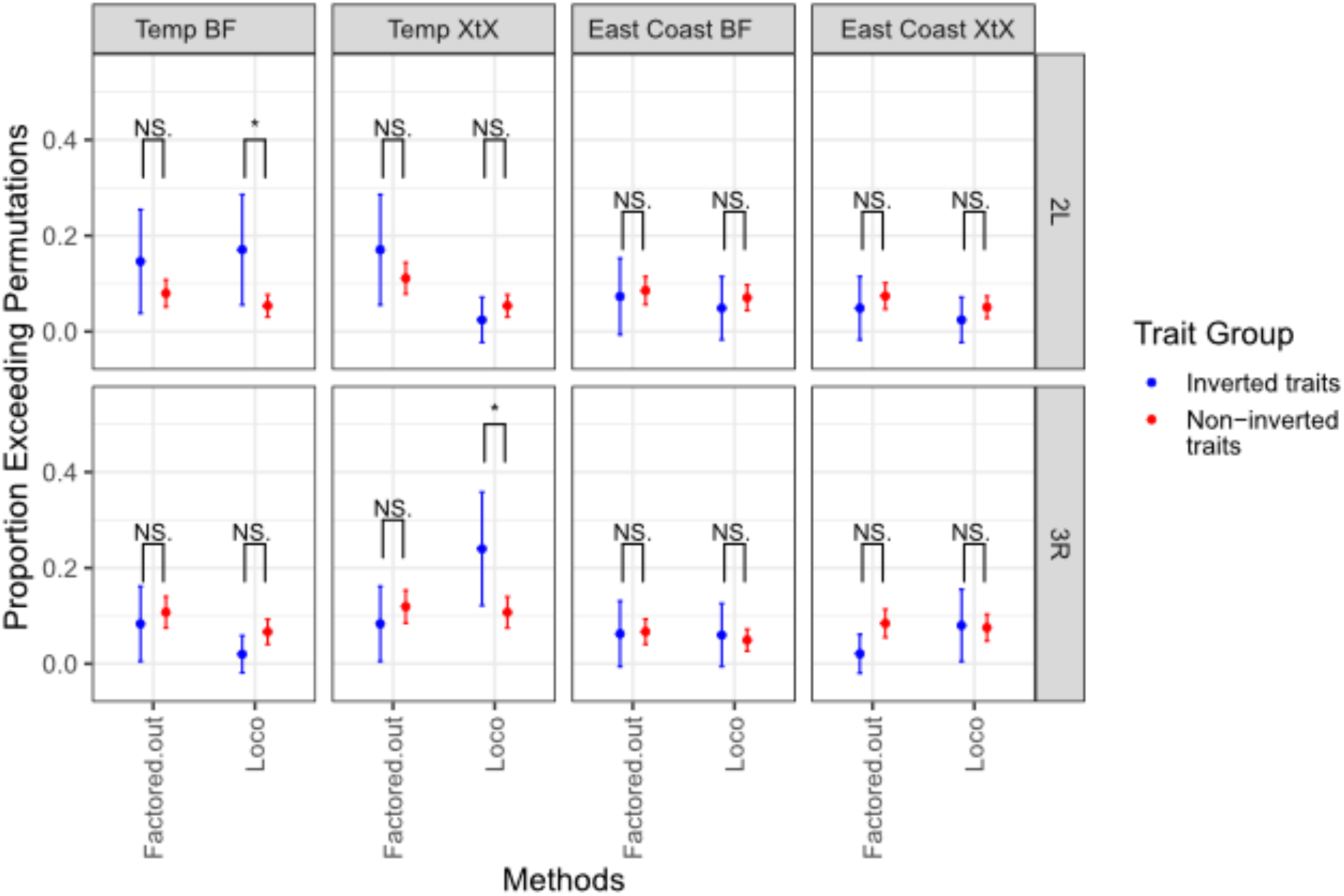
Enrichment of top Bayes loci from two populations with top LOCO and Factored-out GWAS hits. The proportion of traits for whom their enrichment exceeds 95% of permutations is shown with error bars from binomial confidence intervals. Color indicates whether the traits are associated with inversion. (* = p < 0.05, ** = p < 0.01, *** = p < 0.001, NS. = p >= 0.05)

### COLOC enrichment within the genome

To compare the ability of approaches to identify potentially pleiotropic associations of inversions with orthogonal multivariate-traits, we calculated the probability that regions of the genome share polymorphisms that affect multi-dimensional traits (co-localization). Using the top principal component loadings from Fig. 2 as dimensionality reduced traits, we scored effect of SNPs and small indels genome-wide on PC1 and PC2 using the LOCO and Factored-out GWAS approaches. We followed with a sliding window analysis to identify loci within the genome are likely associated with traits differentiated along only PC1, the traits differentiated along only PC2, or for both PC1 and PC2. With the LOCO method, we identified variants within the In(2L)t inverted regions and near the breakpoints that have high association likelihood with PC1 of the In(2L)t-linked traits (**Fig. 6A**) while there was little association on other chromosome arms (Supplemental Figure3). In contrast, for In(3R)Mo areas of likely association were identified across 3R for both PC1 and PC2, with the peaks aligning with other inversion breakpoints on 3R (**Fig. 6B**) and several peaks observed on other chromosomes (Supplemental Figure 4). Notably, the peaks of highest likelihood of association differed between PC1 and PC2, suggesting that distinct loci within the inverted regions influenced different sets of traits. In contrast, SNPs scored using the Factored-out method failed to capture any signal of likely association with either PC1, PC2, or both (Supplemental Figure 5).

**Figure 6.**
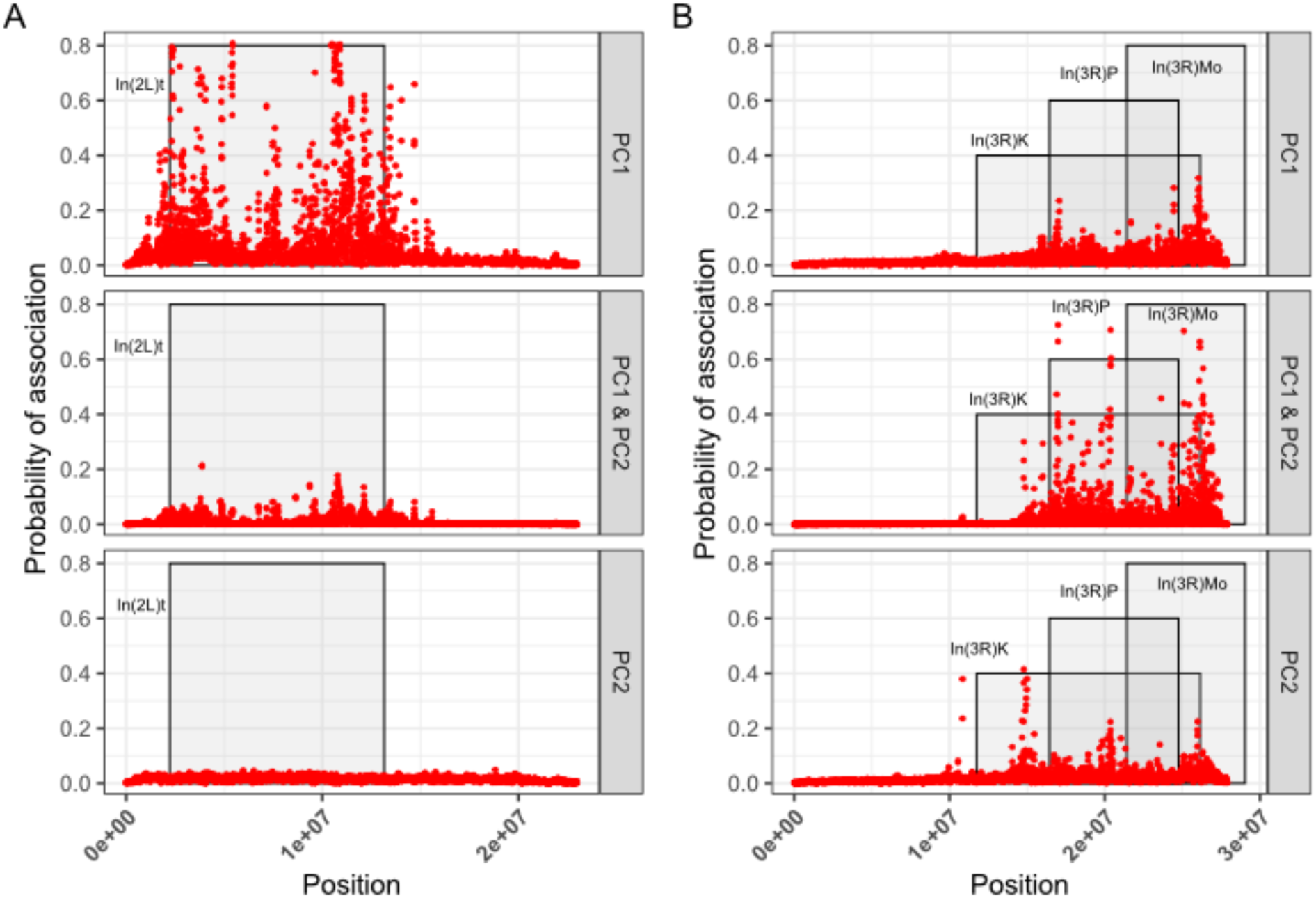
LOCO method reveals areas of likely colocalization of association for inversion-related traits. **A)** Results of a sliding window analysis examining enrichment between SNPs scored using LOCO for PC1 and PC2 of In(2L)t. The y-axis shows the likelihood of association, and the x-axis shows position on the genome. The grey shaded regions show the zone of cosmopolitan inversions on the chromosome arm. **B)** Same analysis as in A, but for traits associated with In(3R)Mo.

## Discussion

Genomic inversions can simultaneously alter multiple traits and provide a mechanism for adaptation. Associations between inversions and traits have been identified across the tree of life, along with evidence for natural selection acting upon these genomic features (6,7,76). We find here that inversions within *D. melanogaster* impact a suite of diverse traits (**Fig. 1A**, **Fig. 2C**, **Fig. 2D**), and specific statistical methods are better equipped to map associations between inversion-linked loci and these traits. Inversions should be considered areas of interest, rather than areas to be skipped over within association studies, as their presence here is shown to explain not only large parts of trait variance (**Fig. 1B**, **Fig. 2B**), but variance within the genome as well (**Fig. 3**). We illustrate that different GWAS approaches have different power in the number and strength of associations they can identify (**Fig. 4**). Compared to several commonly used methods, the LOCO approach is better able to identify variants within inversions associated with key traits, associated with alleles changing across environments (**Fig. 5**), and inversion linked loci that could be pleiotropic (**Fig. 6A**, **Fig. 6B**).

Years of previous work have found numerous insights into the effects of inversion on *D. melanogaster* traits, implicating these mutations in changes to body size, wing size, longevity,, and more (35–41). Despite these known impacts, only 13 out of the 36 publications aggregated here report any test for association between inversions and their trait(s) of study (Supplemental Table 1). Here we reexamined the impact of inversions on a large body of diverse trait data, and show that inversions like In(3R)Mo and In(2L)t significantly affect more traits than would be expected by chance, including inversion-trait associations not previously identified (Supplemental Table 1; **Fig. 1**). In(3R)Mo varies across latitudinal clines in multiple continents (3), and the overlapping inversion In(3R)P is thought to allow for these north-south delineated populations to adapt to different environment (34,36,77,78). In our analysis, In(3R)Mo is likely more enriched than the commonly researched In(3R)P due to its 3x higher presence than In(3R)P within the DGRP.

Here we confirm that inversions on 3R impact body size (**Fig. 2D**). Similarly, highlighting the association between In(2L)t and activity (**Fig. 1A**, **Fig. 2C**) provides new avenues for investigation for the ongoing link to this inversion and seasonal adaptation (43,79). We show that inversion presence explains considerable trait variance within specific traits (**Fig. 1B**, **Fig. 2B**), indicating these inversions should be a major factor in consideration for association studies.

Inversions provide a challenge for association studies, as the increased linkage disequilibrium and relatedness within inverted samples can elevate the false discovery rate (28,74). Many modern GWAS techniques thus seek to compensate for the impact of relatedness by using top principal components as cofactors (80) or factoring our relatedness using GRMs as a random-effect (81). The Factored-out approach described here employs these last two methods, using genome-wide GRMs and additionally factors out the effect of inversions prior to genome-wide association mapping. Of the studies we analyzed, 21/36 used this method or an equivalent for GWA with the DGRP (Supplemental Table 1). However, we report that only about 6% of GWA using this method find more hits than from random permutations (**Fig. 4**), indicating a lack of power and a potentially high false-positive rate amongst many published DGRP studies. Other GWA methods, such as thinning the relatedness matrix for linkage disequilibrium, fare little better (**Fig. 4**). In contrast, LOCO is designed to identify association when there is high LD within the genome, by avoiding proximal contamination between highly linked SNPs while still partially accounting for population structure (56). Recent association studies have using LOCO methods of establishing relatedness while investigating association studies within inversions and other areas of high LD (15,82). For example, Calboli *et al,* 2022 established an association between an agriculturally relevant trout disease and an inversion used a LOCO method, but not with their accompanying “Full” genome method (83). Here we provide evidence that LOCO can outperform other methods at identifying association signal within inverted regions (**Fig. 4**).

Further evidence indicates that the LOCO method provides advantages in identifying loci involved in inversion-mediated adaptation. Recent research indicates that In(2L)t could mediate seasonal adaption (79), and implicated that behavioral traits could be contribute to rapid adaptation to temperature (43,79). By describing the enrichment between our most significant GWAS variants and an independent set of environmentally relevant alleles, we illustrate within In(2L)t alleles identified as both highly temperature associated and as top GWAS hits for traits such as sleep and startle response. (**Fig. 5**). The overlap of environmentally-varying alleles and top GWAS loci can be identified from LOCO-based GWAS, but not from the Factored-out approach (**Fig. 5**).

We identify differences in the ability of GWAS approaches to identify regions within the genome that are associated with multiple traits (pleiotropy). The ability of inversions to pleiotropically effect multiple traits has already been noted in salmonids (83,84), and mice (6,85). Dimensionality reduction of multiple traits can potentially indicate some shared mechanism or biological component that explains variance in aspects of morphology (86,87). For example, different aspects of body size can load onto the top principal component, reflecting some unifying aspect of body size development (88). In contrast, phenotypic variation across principal components can reflect a different degree of pleiotropy across orthogonal traits. Thus, we characterized areas of high association between PC1 and PC2 of the inversion linked trait sets to illustrate such pleiotropy using a colocalization test. Only the LOCO approach identifies that the areas of highest association with multiple inversion-linked traits are near their corresponding inversions, as one might expect (**Fig. 6A**, **Fig. 6B**). However within the inverted regions there are peaks of higher likelihood of association, similar to the finding in Nunez et al., 2024 of peaks of SNP-phenotype enrichment within In(2L)t (43). Peaks of association with PC1 of In(3R)Mo may indicate loci relevant to traits such as body size, while a peak for PC2 may indicate different loci relative to traits such as metabolic storage and sleep, and the peaks for both PCs indicate areas of likely pleiotropic effect (**Fig. 2D**). While LOCO identifies these areas of likely association, this signal cannot be recapitulated using the Factored-out approach (Supplemental Figure 5).

Inversions have the potential to be a fruitful area of investigation within association studies rather than a statistical nuisance. Despite the evidence across taxa that inversions can alter many classes of traits (6,7,76), inversions are sometimes presented as a statistical hindrance (28,74). Efforts such as the creation of popular mapping populations from largely co-linear genotypes (23–25), and the use of multiple methods to factor out inversion presence within the DGRP (26) represent steps to account for these mutations. To be clear, the phenotyping and association studies from the DGRP and other mapping populations have produced many important and foundational insights. However, models that remove or ignore inversions miss a valuable opportunity. Methods like LOCO offer tools toward building association studies to identify relevant loci within inverted regions (82,83). Inversions can play a significant role in the traits of humans and across many forms of life (89–92). Improving our ability to connect inversion to traits will motivate future work to better understand how these complex mutations contribute to trait regulation and formation.

## Data Availability

The DGRP’s genomic data, Wolbachia infection status, “Full” GRM, and inversions genotype table are all available from the DGRP website (http://dgrp2.gnets.ncsu.edu/data.html). The aggregated DGRP phenotype data is available from the DGRPPool website (https://dgrpool.epfl.ch/) or from their original publications. All other data is available for download at (https://github.com/benedictlenhart/InversionGWAS)

## Acknowledgements

We thank Research Computing at UVA for the use of computational resources, and for the staff’s patient and consistent support (https://rc.virginia.edu), and John Teurman for his work annotating information from *Drosophila* publications.

