## Supplementary Figures for "Cosmopolitan inversions have a major impact on trait variation and the power of different GWAS approaches to identify associations"

### Supplemental Figures

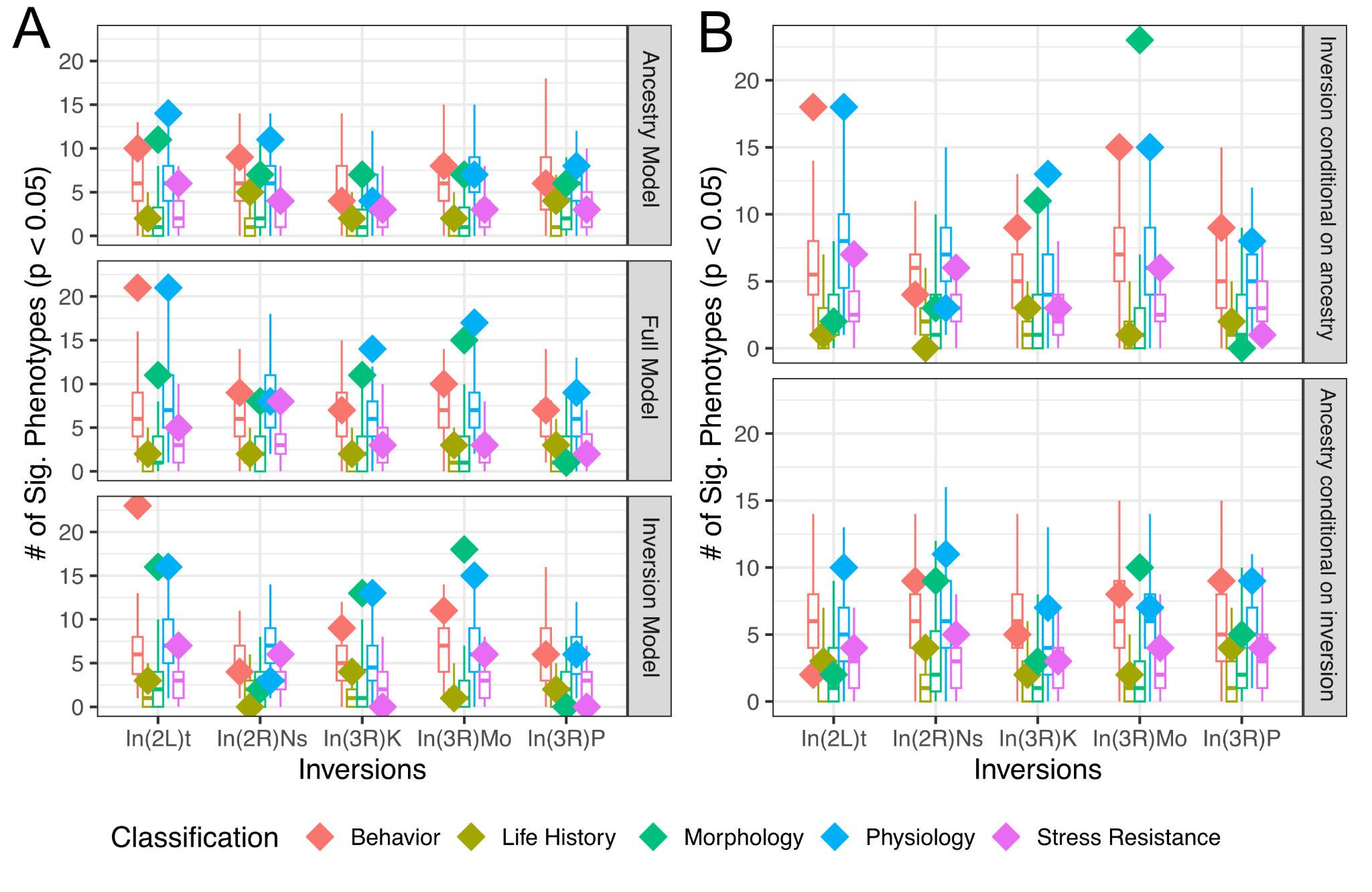

**Supplemental Figure 1: The addition of ancestry does not remove the broad impact of inversion genotype on phenotype.** A The number of phenotypes with significant associations is shown as diamonds for the Ancestry and Inversion model, as well as for a Full model that uses both ancestry and inversion genotype as fixed effect. A set of paired 100 permutations of each model is shown as a box and whisker plot. Results are split across five cosmopolitan inversions, and colored by trait classification. B The same plot as in A, now showing a comparison between the Full and Ancestry models, as well as the Full and Inversion models.

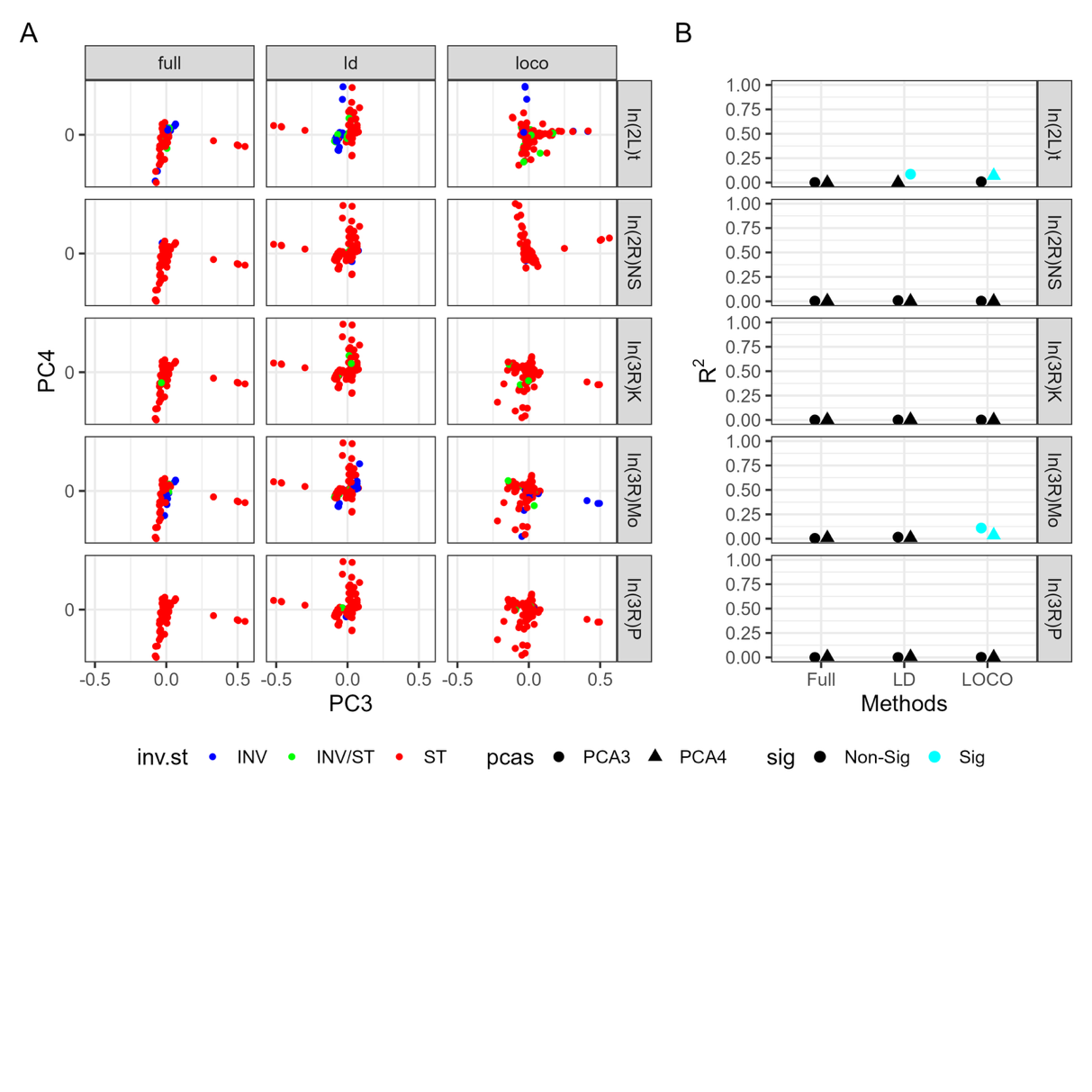

**Supplemental Figure 2:** Genomic principal components PC3 and PC4 have little correlation with inversion genotype. **A)** The third and fourth genomic PCs for each sample colored by the genotype of that sample. **B)** The R^2^ values for models comparing PC3 and PC4 to inversion, colored by which values exceed a distribution of permutations.

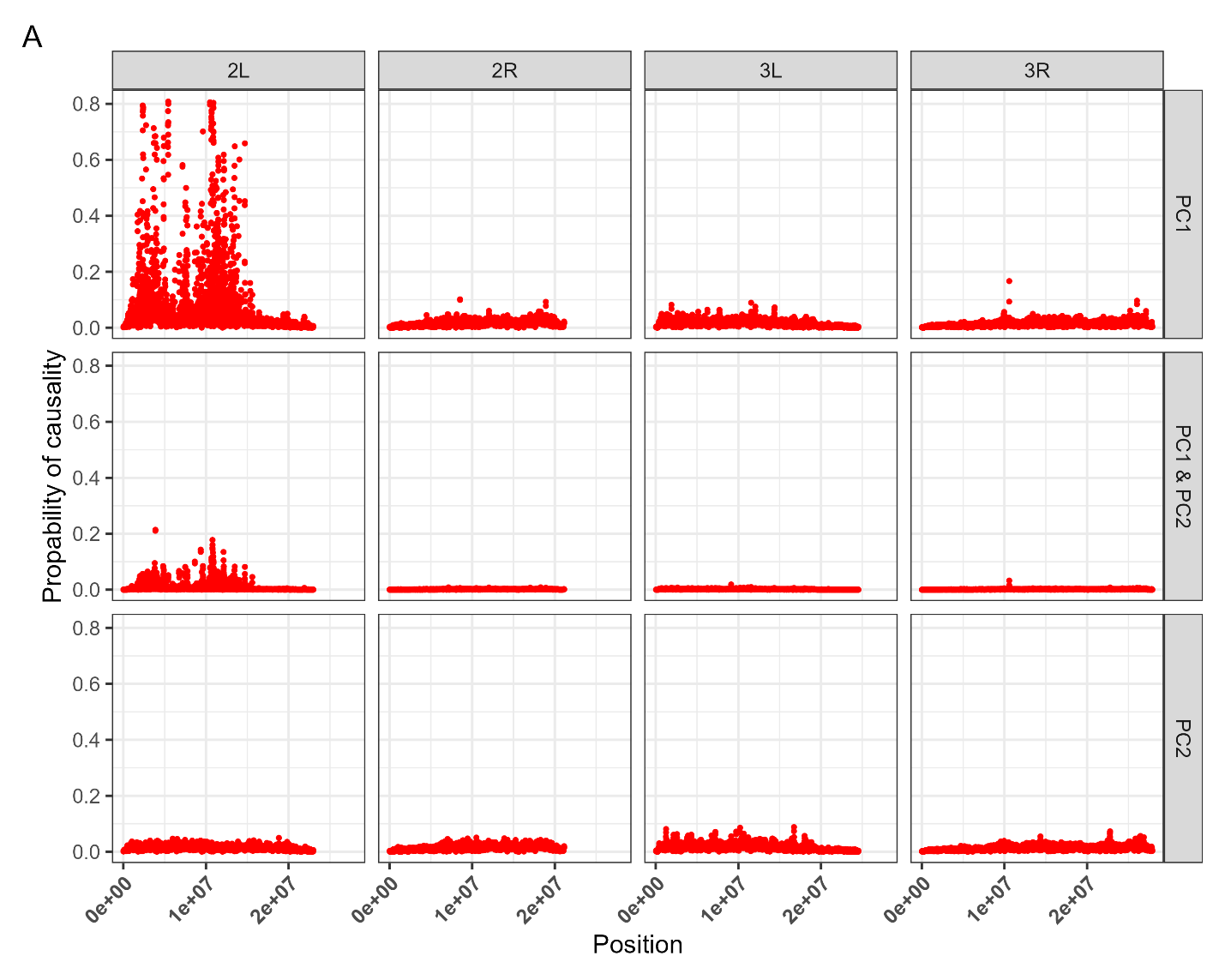

**Supplemental Figure 3.** Signal of loci association with In(2L)t is mostly adjacent to the inversion. The same results of the association study using the LOCO method from Fig 6 are shown across the genome, showing the likelihood of a SNP’s association with PC1, PC2, or both from the In(2L)t PCA analysis

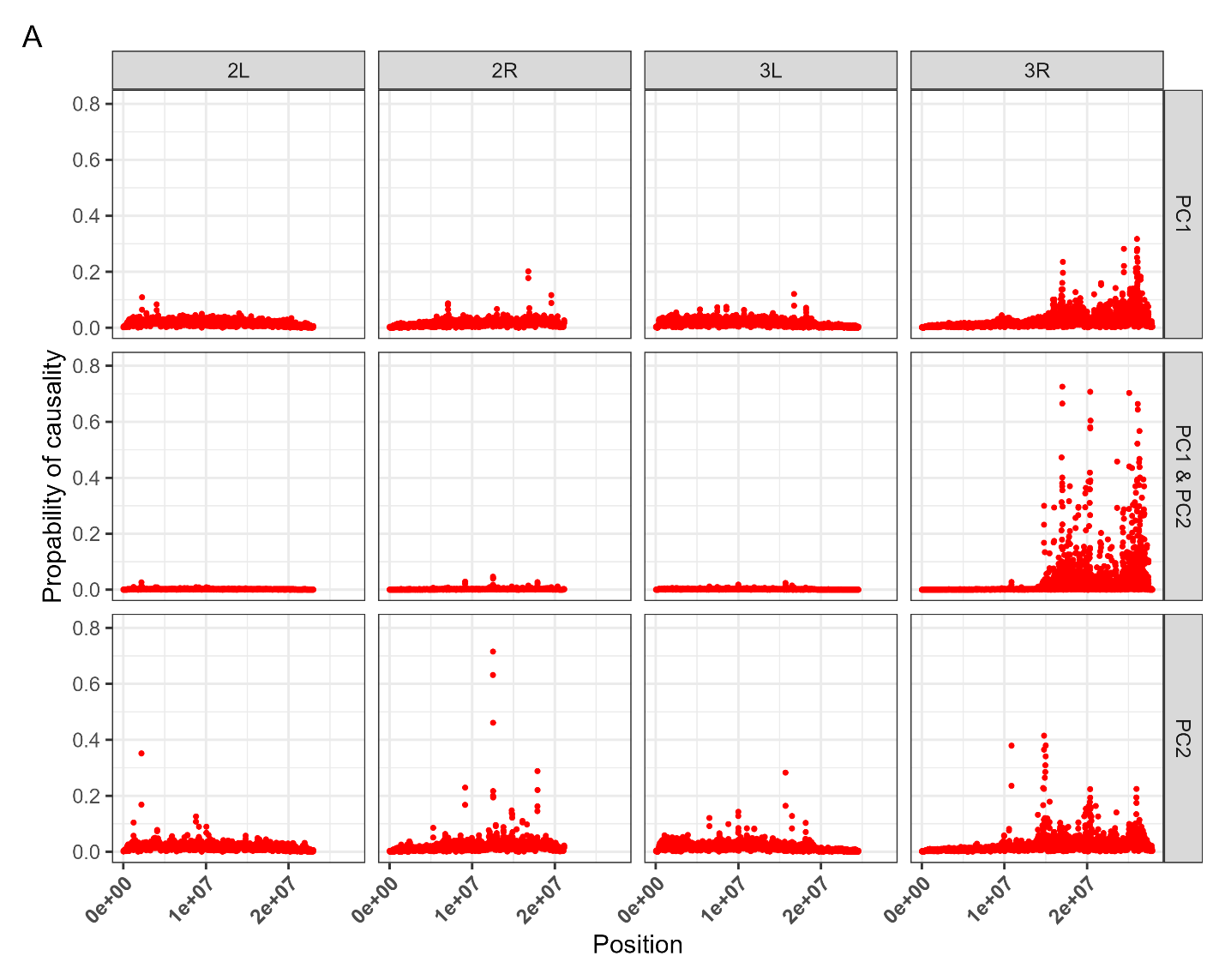

**Supplemental Figure 4.** Signal of loci association with In(3R)Mo is elevated on 3R. The same results of the association study using the LOCO method from Figure 6 are shown across the genome, showing the likelihood of a SNP’s association with PC1, PC2, or both from the In(3R)Mo PCA analysis

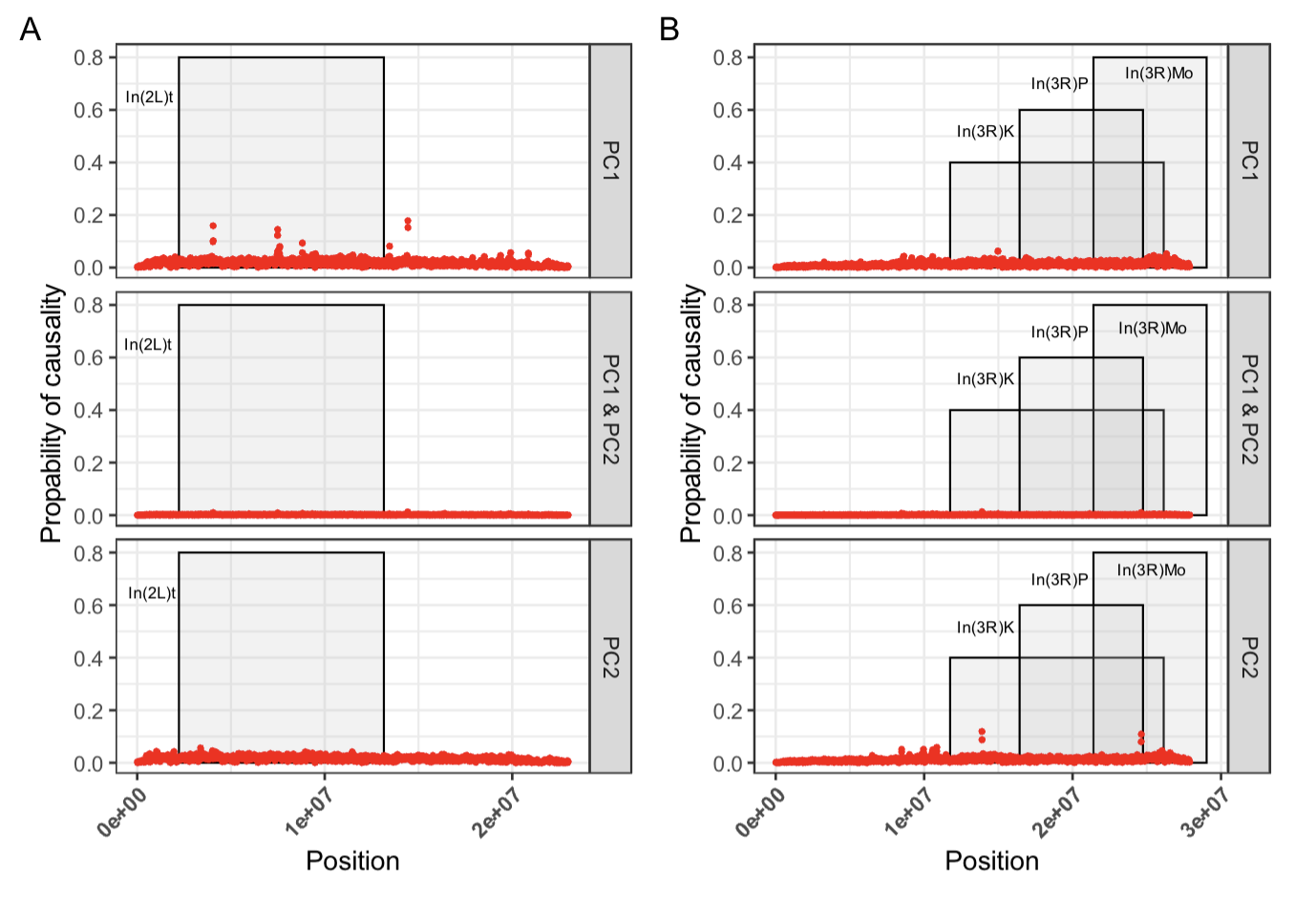

**Supplemental Figure 5.** Factored-out method fails to identify areas of likely association. **A)** Results of a sliding window analysis examining enrichment between SNPs on 2L scored using Factored-out for PC1 and PC2 of In(2L)t, the y- axis shows the strength of enrichment and the x-axis shows position on the genome. Grey shaded region show the zone of cosmopolitan inversions on the chromosome arm. **B)** Same analysis as in A, but considering chromosome arm 3R and inversion In(3R)Mo.
